# 2dSS: a web server for protein secondary structure visualization

**DOI:** 10.1101/649426

**Authors:** Diksha Priya Lotun, Charlotte Cochard, Fabio R.J Vieira, Juliana Silva Bernardes

**Affiliations:** Sorbonne Universités, UFR 919, 75005 Paris, France; Institut de Biologie de lÉcole Normale Supérieure (IBENS), 75005 Paris, France; Sorbonne Universités, laboratoire de Biologie Computationnelle et Quantitative, CNRS UMR7238, 75005 Paris, France

**Keywords:** secondary structure prediction, multiple sequence alignment, homologous proteins

## Abstract

2dSS is a web-server for visualising and comparing secondary structure predictions. It provides two main functionalities: 2D-alignment and compare predictions. The “2D-alignment” has been designed to visualise conserved secondary structure elements in a multiple sequence alignment (MSA). From this we can study the secondary structure content of homologous proteins (a protein family) and highlight its structural patterns. The “compare predictions” has been designed to compare the output of several secondary structure prediction tools, and check their accuracy when compared with real secondary structure elements extracted from 3D-structure. 2dSS provides a comprehensive representation of protein secondary structure elements, and it can be used to visualise and compare secondary structures of any prediction tool.

**Availability:** http://genome.lcqb.upmc.fr/2dss/

## Introduction

The secondary structure (SS) of proteins is a useful resource for three-dimensional (3D) structure prediction, fold-recognition and protein classification. SS refers to the local conformation of the polypeptide chain of proteins. Each amino acid in a protein is associated to a secondary structure element (SSE). There are two regular SSEs: *α*-helix and *β*-sheets, and one irregular: coil region, that connects regular elements. A number of methods have proposed for predicting SSEs in protein sequences (1–4), and each tool has its owner webserver with a particular visualisation system. Most of existent tools provide SSE predictions for a single sequence at time, or for a set of unaligned sequences. However, it could be very useful: (i) to visualise the SS conservation in a multiple sequence alignment (MSA) for obtaining a consensus overview in a given protein family, (ii) to compare the SS predictions of different tools for visualising (dis)agreement regions, and (iii) to check the accuracy of SS tools by comparing their predictions to the real SSEs extracted from 3D-structure.

Recently, MPI bioinformatics Toolkit proposed two tools: Ali2D and Quick2D (5) that address the points (i) and (ii) respectively. Ali2D allows users to visualise SS conservation in a MSA, where SSE predictions of each sequence in the MSA are produced by PSIPRED (2), and Quick2D compares the SSE predictions of four tools: PSIPRED, SPIDER2 (1), PSSPred (3) and DeepCNF-SS (4). Although Ali2D and Quick2D have become important resources for SS prediction, their visualisation system is very simple and both are limited to a fixed number of prediction tools. Moreover, these tools do not address the point (iii) of not allowing comparisons with the 3D-structure. Here, we propose 2dSS, a SS visualization web server, which addresses the three relevant points previously mentioned. 2dSS adopts the standard format for SSEs based on the three-state codes: H for *α*-helix, E *β*-sheets, and C for coil regions. Since these three-state codes can be derived from the output of SS tools, 2dSS works for any tool. Moreover, 2dSS does not run prediction tools, but import their outputs, being extremely fast and providing an easy and interpretable picture that can be downloaded in many formats. 2dSS also accepts the Ali2D and Quick2D outputs by producing a more user-friendly graphical representation for SSEs.

## Web interface

The web-server has two main functionalities: “view 2D-alignment” that addresses the item (i), that is, it allows users to visualise and compare SSEs in a set of aligned sequences (a protein family), and “compare predictions” that addresses items (ii) and (iii) by comparing SS predictions of several tools and allowing to check the prediction accuracy. In any case, users can set colors for SSEs and save their pictures in SVG format, for a posterior edition, or in other image format such as PNG, and PDF. In the next sections, we detail the two main functionalities, and more details and examples can be found in the Help tab of 2dSS website.

### View 2D-alignment

Given a MSA, representing a protein family, and the predicted SSEs of its constituent sequences, we can map each SSE onto a MSA position to obtain a 2D-alignment. In 2dSS, the user provides a MSA in Fasta or Clustalw format and the SSE predictions of each aligned sequence, to obtain a 2D-alignment that shows the SS conservation for that MSA. The 2D-alignment highlights the secondary structure conservation in a set of related sequences, where is possible to see how SSEs of homologous proteins are aligned. In particular, it can be used to compare structurally homologous proteins with low sequence identity and detect their SS patterns. For instance, Figure 1-a shows the partial 2D-alignment of five sequences of Microbial ribonucleases superfamily (SCOP (6) accession number d.1.1). Proteins in this superfamily have less than 30% of sequence identity, but we can still observe SSE conservation among different protein sequences. The Microbial ribonucleases superfamily is composed by three families: Fungal ribonucleases (d.1.1.4), Ribotoxin (d.1.1.3) and Bacterial ribonucleases (d.1.1.2). From Figure 1-a, we note that SSEs of Fungal ribonucleases sequences are closer to SSEs of Ribotoxin, and slightly different from Bacterial ribonucleases ones. It is expected since Fungal ribonucleases and Ribotoxin families are constituted by fungal sequences, while Bacterial ribonucleases by bacteria sequences. Note that 2dSS does not perform MSA or run SSE prediction tools, but it provides the possibility of uploading Ali2D output for a better visualization.

**Fig. 1.**
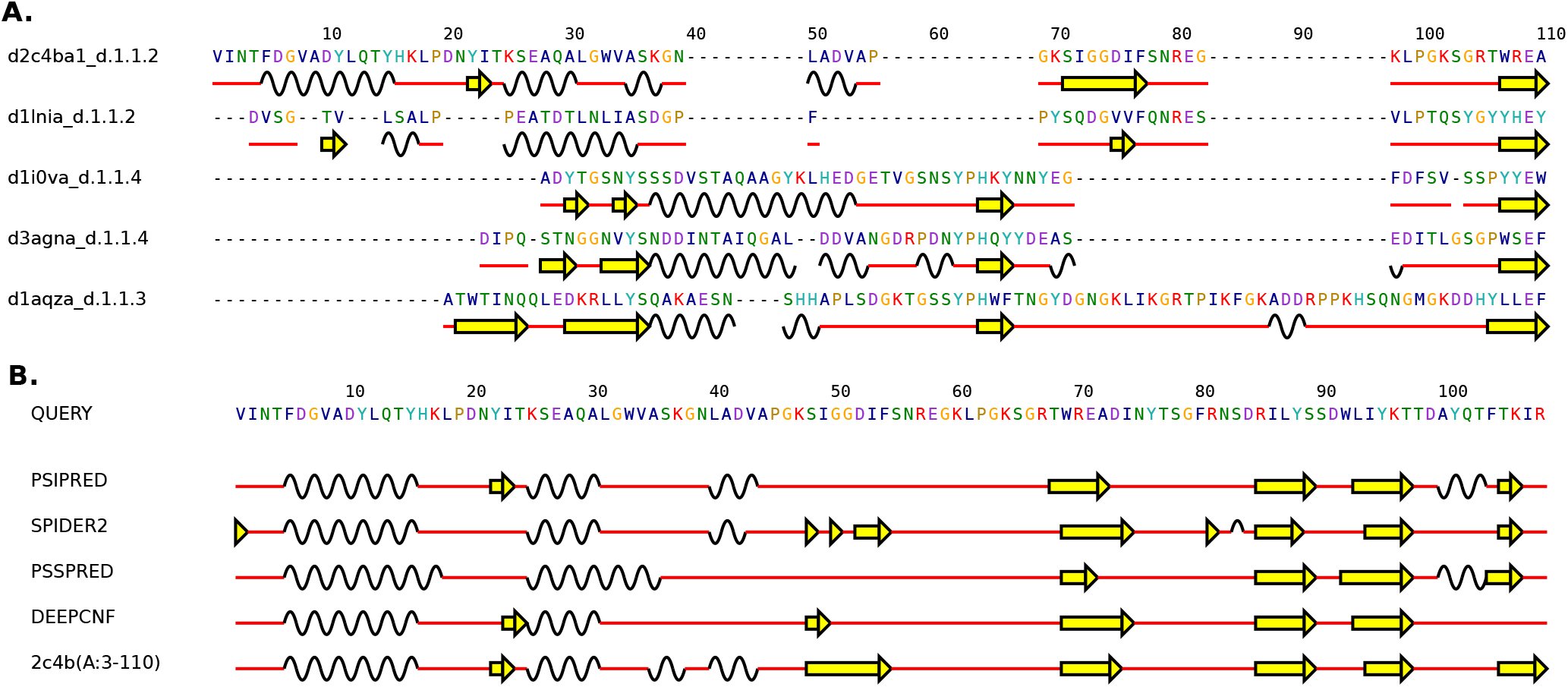
The two main functionality of 2dSS: view 2D-alignment (A) and compare predictions (B)

### Compare predictions

The functionality “Compare predictions” was designed to identify differences in SSE assignments among a set of prediction tools, and eventually compare those assignments to the real SSEs extracted from the 3D structure. For that, the user provides a protein sequence, SSEs assigned by different prediction tools, and optionally the real SSEs extracted from the 3D-structure, that is, from the PDB file (7). Note that 2dSS can extract the real SSEs automatically. For that, the user must provide the DSSP file obtained in http://www.cmbi.ru.nl/xssp/ and specify the chain, start and end amino acid positions. Figure 1-b shows SSE comparisons for Inhibitor cystine knot protein (PDB accession number 2c4b, chain A:3-110). We start by showing the amino acid sequence of Inhibitor cystine knot protein, follow by the SSE predictions obtained by four different tools, and the real SSEs extracted from PDB file. We can observe some differences among predictions obtained by the four investigated tools. SPIDER2 and PSSPRED were not able to predict the first *β*-sheet localised at position 22, while DeepCNF seems to be more accurate, predicting correctly most of SSEs. By comparing SS predictions, we can pint point prediction errors of some tools, and identify regions in the protein where SSE prediction is harder. Note that 2dSS is not able to run SS prediction tools, but it provides the possibility of uploading Quick2D output file. Quick2D allows to run four SS prediction tools, simultaneously.

## Comparison with other tools

We compare 2dSS to Ali2D and Quick2D, both tools belongs to MPI bioinformatics Toolkit. Figure 2 shows the 2D-alignment of Microbial ribonucleases superfamily produced by 2dSS (Fig. 2-a) and by Ali2D (Sup 2-b). Since 2dSS uses graphical elements such as arrows and waves, it provides a clearer visualisation of SSEs than flat text proposed by Ali2D. Moreover, 2dSS can separate SSEs from sequence alignment to have a better view of secondary structure content, as shown in Fig. 3. The output comparison between 2dSS and Quick2D are shown in Fig. 4-ab, respectively. For this comparison, we used the Inhibitor cystine knot protein sequence, shown in Figure 1-b, and four prediction tools embedded on Quick2D. Note that 2dSS provides a more friendly visualisation, but also allow user to compare predictions to the real SSEs extracted from PDB. Another advantage of 2dSS compared to Ali2D and Quick2D is the possibility of setting results with different colors, and save pictures in SVG that are editable, or choose among PNG or PDF format.

**Fig. 2.**
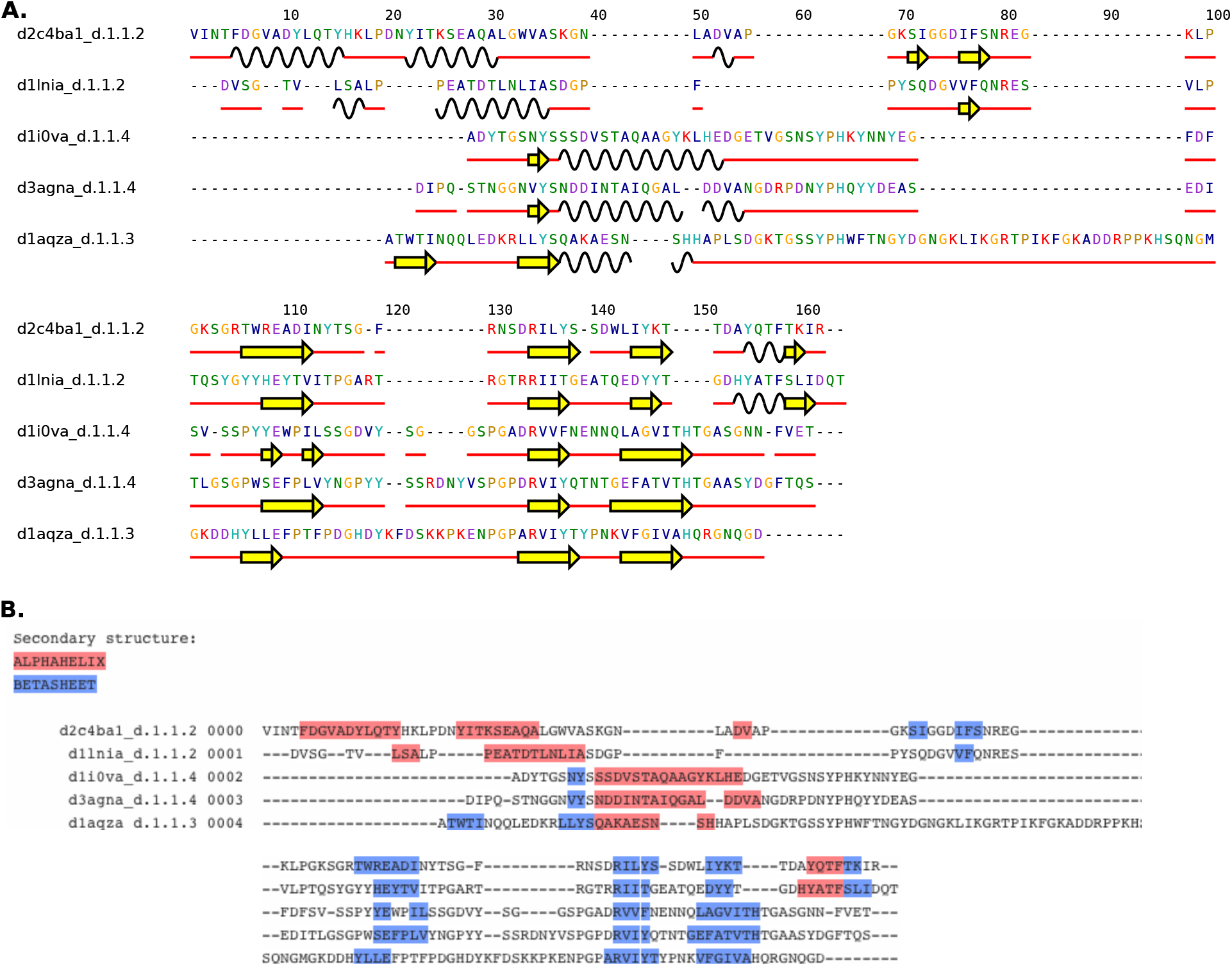
Comparing 2D-alignment outputs of 2dSS (A) to Ali2D (B). The sequences of Microbial ribonucleases superfamily (SCOP accession number d.1.1) were aligned with clustal Omega and inputted in the Ali2d web server that uses PSIPRED to predict the secondary structure elements of each aligned sequence. The output of Ali2D is a flat text where alpha-helix, beta-sheets and coils are showed in red, blue and black, respectively. To produce 2dSS output, we used a not formatted output of Ali2D. 2dSS webserver uses graphical elements to represent *α*-helices (black waves), *β*-sheet (yellow arrows), and coil (red straight lines)

**Fig. 3.**
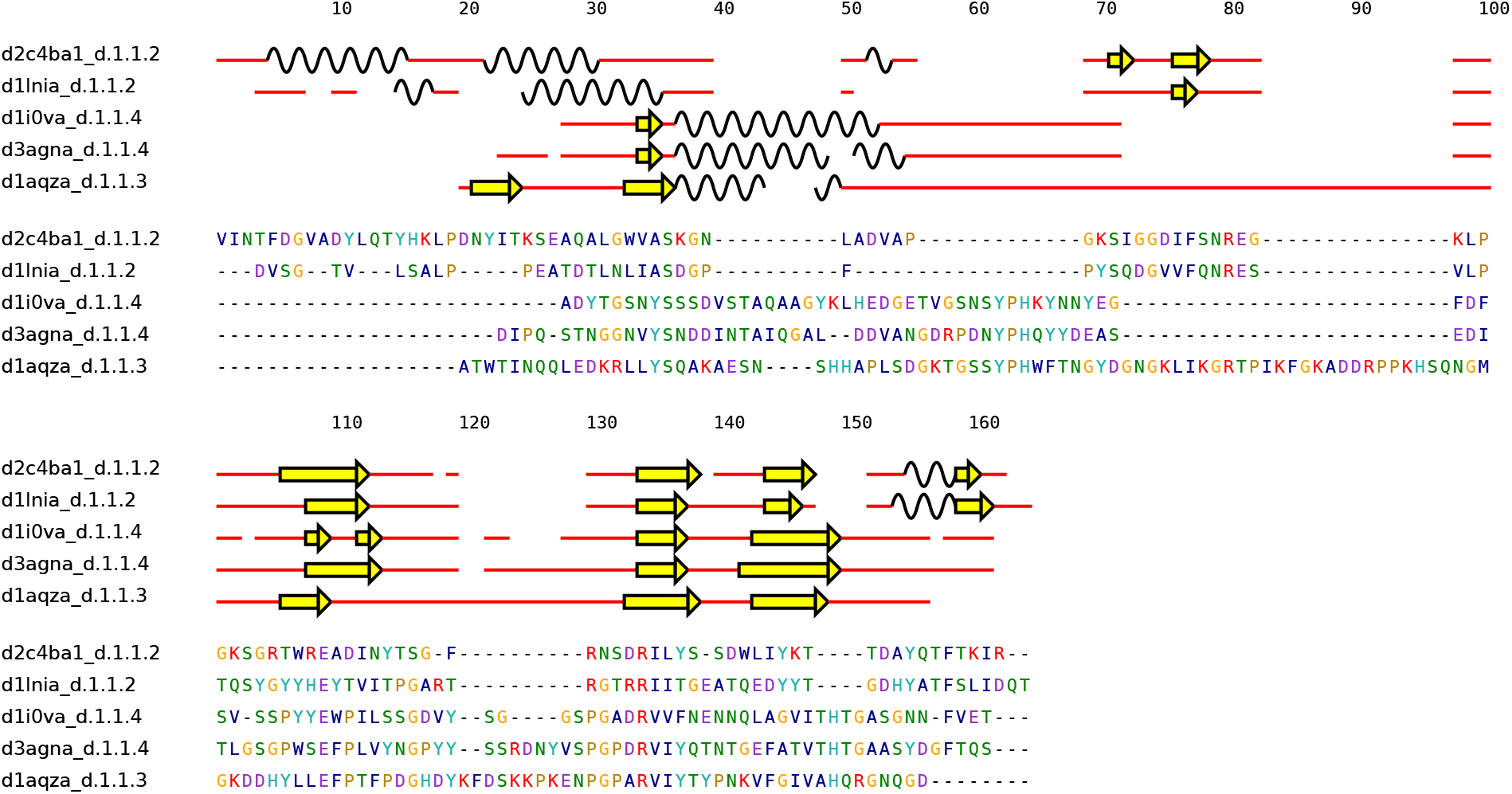
Separated views of 2D-alignment. By checking the option separate in “view 2d-aligment” menu of 2dSS webserver, uses can separate the secondary structure elements from the Multiple sequence alignment.

**Fig. 4.**
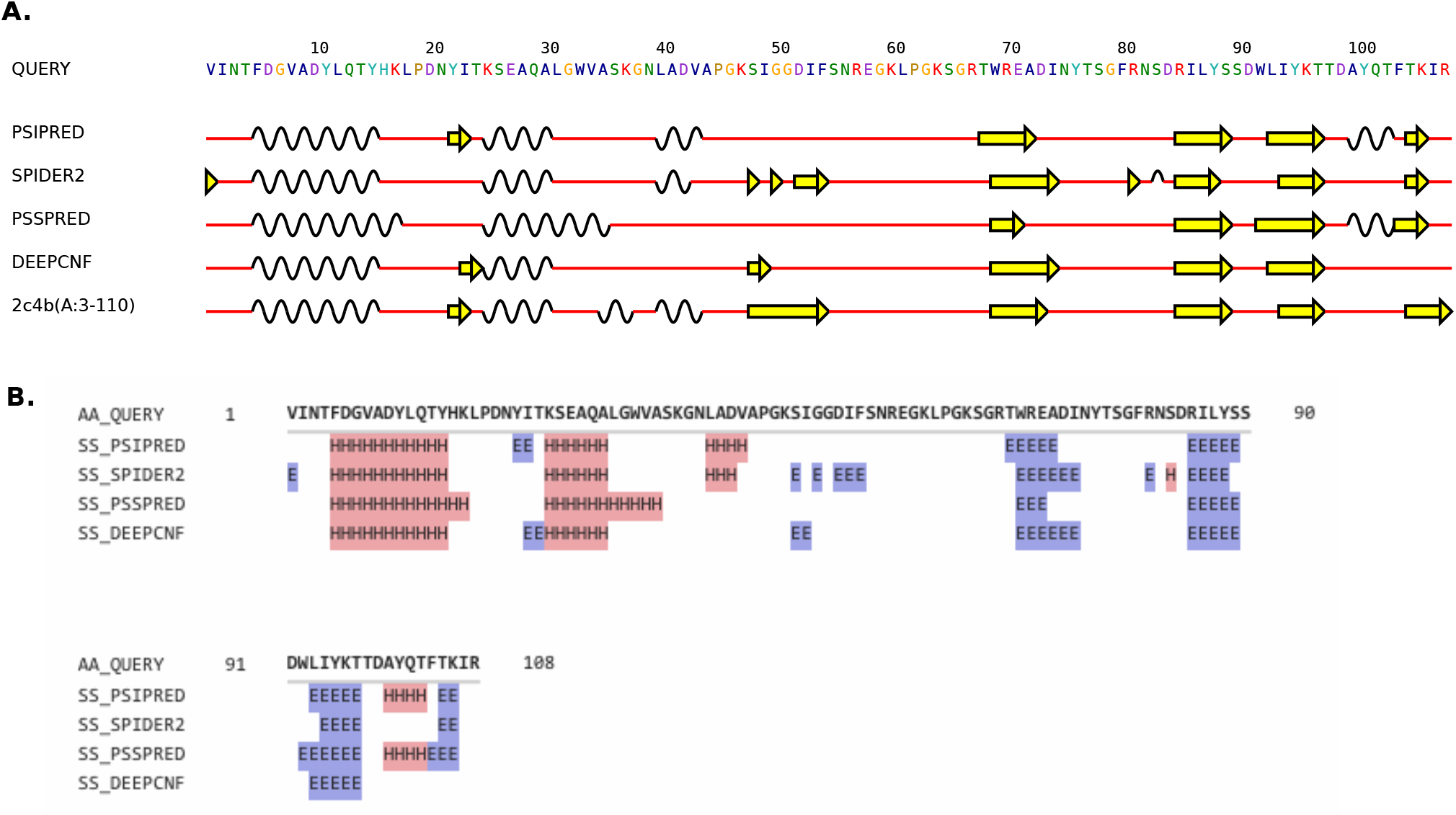
Comparing “compare predictions” outputs of 2dSS (A) to Quick2D (B). We extracted the amino acid sequence of Inhibitor cystine knot protein from the PDB file (Accession number 2c4b, chain A:3-110). We input this sequence in Quick2D to obtain the secondary structure predictions of PSIPRED, SPIDER2, PSSPred and DeepCNFSS. The output of Quick2D is a flat text where *α*-helices, *β*-sheets and coils are showed in red, blue and white, respectively. To produce 2dSS output, we input the Quick2D output in the 2dSS webserver. In order to generate the real secondary structure elements of Inhibitor cystine knot protein, we also input the DSSP file of 2c4b. 2dSS output shows graphical elements that represent *α*-helices (black waves), *β*-sheets (yellow arrows), and coils (red straight lines). Moreover, it shows the real secondary structure elements that allow users to check the accuracy of prediction tools.

## Conclusion

2dSS is a unique web server for secondary structure visualisation. It is simple and fast, since it does not run itself SSE prediction tools. Instead, it accepts the output of main prediction tools and formats them in a friendly and editable format. 2dSS can be used for a multitude of purpose: as a means of determining the secondary structure content of a protein family; for visual display of differences between homologous proteins or wild-type, and mutant protein structures; for comparing the SSEs of protein structures prepared under different crystallization conditions; and to enable comparisons of structurally homologous proteins and determine how closely their SSE assignments are.

